# Identification of Scaffold Specific Energy Transfer Networks in the Enthalpic Activation of Orotidine 5’-Monophosphate Decarboxylase

**DOI:** 10.1101/2025.01.29.635545

**Authors:** Pankaj Dubey, Anish Somani, Jessica Lin, Anthony T. Iavarone, Judith P. Klinman

## Abstract

Orotidine 5’-monophosphate decarboxylase (OMPDC) is one of the most efficient enzyme systems studied, enhancing the decarboxylation of OMP to uridine 5’-monophosphate (UMP) by ca. 17 orders of magnitude, primarily by reducing the enthalpy of activation by ca. 28 kcal/mol. Despite a substantial reduction in activation enthalpy, OMPDC requires 15 kcal/mol of activation energy post- ES complex formation. This study investigates the physical basis of how thermal energy from solvent collisions is directed into the active site of enzyme to enable efficient thermal activation of the reaction. Comparative study of temperature-dependent hydrogen-deuterium exchange mass spectrometry (TDHDX) for WT and mutant forms of enzymes has recently been shown to uncover site specific protein networks for thermal energy transfer from solvent to enzyme active sites. In this study, we interrogate region-specific changes in the enthalpic barrier for local protein flexibility using a native OMPDC from *Methanothermobacter thermautotrophicus* (Mt-OMPDC) and a single site variant (Leu123Ala) that alters the activation enthalpy for catalytic turnover. The data obtained implicate four spatially resolved, thermally sensitive networks that originate at different protein/solvent interfaces and terminate at sites surrounding the substrate near the substrate phosphate-binding region (R203), the substrate- ribose binding region (K42), and a reaction enhancing loop5 (S127). These are proposed to act synergistically, transiently optimizing the position and electrostatics of the reactive carboxylate of the substrate to facilitate activated complex formation. The uncovered complexity of thermal activation networks in Mt-OMPDC distinguishes this enzyme from other members of the TIM barrel family previously investigated by TDHDX. The new findings extend the essential role of protein scaffold dynamics in orchestrating enzyme activity, with broad implications for the design of highly efficient biocatalysts.

## Introduction

Orotidine 5’-monophosphate decarboxylase (OMPDC) is an extraordinarily efficient enzyme that accelerates the unimolecular decarboxylation of Orotidine 5’-monophosphate (OMP) to Uridine 5’- monophosphate (UMP) by an average of 17 orders of magnitude (*k*_cat_*/k*_non_,), corresponding to a reduction in the free energy barrier for decarboxylation of ca. 22 kcal/mol.^1,2^ This substantial reduction is largely due to a decrease in the observed activation enthalpy (Δ*H*^‡^), with minimal contribution from entropy changes (TΔ*S*^‡^), cf. Fig. 1a.^3,4,5^ The majority of other highly efficient enzymes^6,7^ also achieve their catalytic power by significantly lowering their activation enthalpies, underscoring the crucial role of reduced enthalpic barriers to effect high catalytic rate enhancements. OMPDC functions exclusively in its dimeric form, with all known crystal structures exhibiting a homo-dimeric TIM barrel fold.^8,9^ A comparison of crystal structures for the apo-form of *Methanothermobacter thermautotrophicus* (Mt- OMPDC) to the enzyme bound with 6-azaUMP , a tight-binding competitive inhibitor described as a transition state analog (TSA),^10, 11^ reveals a regional conformational change (RMSD of 0.5 Å) that includes a loop7 closure and shielding of the active site from solvent exposure, Fig. 1b. The active site of Mt-OMPDC in the presence of 6-azaUMP further indicates the importance of a tetrad of charged residues, a phosphate-binding residue (R203), and with a cluster of hydrophobic residues within the barrel cavity (Fig. 1c).

**Figure 1.**
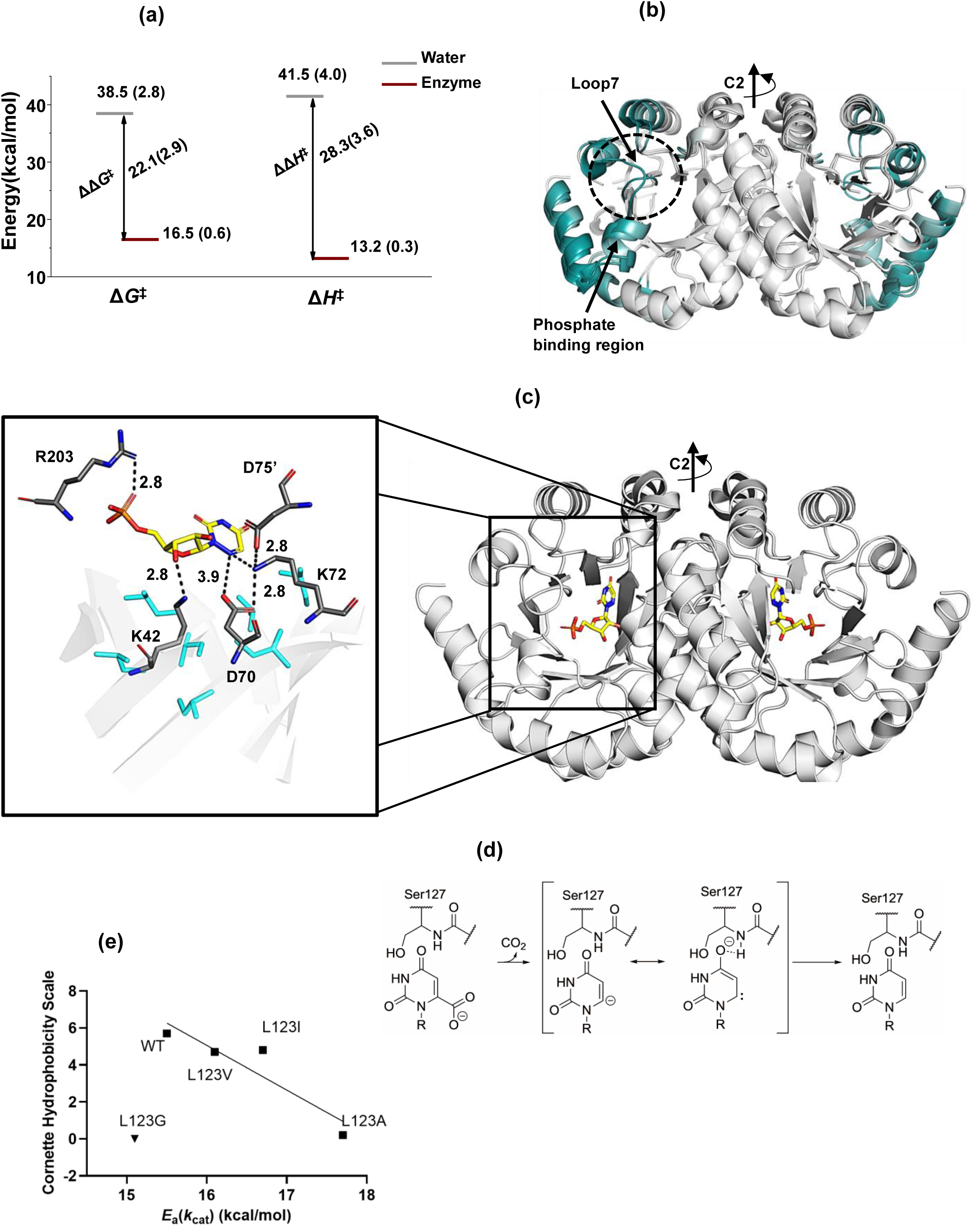
Structural and Energetic Insights into OMP Decarboxylation Catalyzed by OMP Decarboxylases. (a) Free energy (Δ*G*^‡^) and activation enthalpy (Δ*H*^‡^) for OMP decarboxylation, showing the energy differences between the reaction in water and in the presence of enzyme. Data for the reaction in water were derived from OMP analogue decarboxylation in aqueous solution,^4^ followed by averaging, while enzyme data were based on the averaged reaction energetics of *Escherichia coli* OMP decarboxylase (Ec*-*OMPDC), *Saccharomyces cerevisiae* OMP decarboxylase (Sc-OMPDC), and *Methanothermobacter thermautotrophicus* OMP decarboxylase (Mt-OMPDC) under *k*cat conditions.^5^ (b) Crystal structure overlay of Mt-OMPDC in its open (apo) form (PDB: 3G18) and ligand-bound (6-azaUMP) closed form (PDB: 3G1A). Residue-specific RMSD values were calculated to identify structural changes upon ligand binding and visualized on the crystal structure using the PyMOL script ColorbyRMSD,^21^ with a white-gray-deepteal color gradient. Regions showing minimal structural differences are depicted in white, while regions with increasing RMSD values are colored deep teal. Loop7 (residues 180-188), highlighted by a dotted circle, is unstructured in the apo form and adopts a well-ordered, closed conformation over the ligand in the bound form. Due to its unstructured nature in the apo form, RMSD values for this loop could not be calculated, and it was manually colored deep teal for visual clarity. (c) Crystal structure of the Mt-OMPDC homodimer bound to the tight binding analogue 6-azaUMP (PDB: 3G1A), where 6-azaUMP is shown in yellow stick representation. The active site of Mt-OMPDC is highlighted, featuring the catalytic tetrad (K42, D70, K72, and D75’) along with R203, shown in dark gray with labeled interaction distances between the substrate and these residues. Hydrophobic residues from the β-strands (I16, I68, I96, L123, V155, I178, I200), which project into the barrel, are depicted in aqua stick representation but are not labelled. These hydrophobic residues form van der Waals contacts with neighboring hydrophobic residues. (d) Mechanism of OMP decarboxylation through the formation of a vinyl carbanion. (e) Linear relationship between the Cornette hydrophobicity scale ^22^ of the residue at position 123 of Mt-OMPDC and the activation energy (*E*a(*k*cat)) for the reaction (R² = 0.85), excluding the L123G mutant, which is represented as an inverse triangle and shown for comparison only.

The consensus mechanism for OMP decarboxylation involves a rate-limiting formation of a vinyl carbanion-carbene intermediate, followed by protonation from residue K72 (Fig. 1d).^12–16^ A pK_a_ analysis of the C-6 proton of enzyme bound product UMP shows a 10-unit reduction when compared to water, consistent with enhanced stabilization of a carbanion-carbene intermediate.^13^ While the proximal K72 moiety may contribute to electrostatic stabilization, D70 and D75’ groups will introduce destabilization. Interestingly, the K72A mutation reduces specific activity by about five orders of magnitude but also significantly lowers the dissociation constant (*K*_d_) for UMP by the same extent, indicating that the reduced activity is likely due to product inhibition rather than a loss of transition state stabilization.^17^ One of the most intriguing aspects of the active site is its lack of functional groups that typically would stabilize a local carbanion intermediate. This has led to the proposed carbene intermediate that is stabilized by H-bonding from a remote S127 backbone amide in loop 5. However, modifying the amide group of S127 to an ester resulted in only a two order-of-magnitude decrease in *k*_cat_, indicating that this single interaction contributes only a small fraction to the overall catalytic enhancement.^18^

Substrate destabilization by steric crowding and electrostatic repulsion from the D70 side-chain has also been proposed to produce an out-of-plane bending of the departing CO_2_ group,^10,19^ supported by binding studies of 6-CN-UMP and 6-N_3_-UMP.^20^ However, mutational and structural analyses again conclude that such substrate destabilization will contribute only about 3-4 kcal/mol to the total catalytic enhancement.^10^ Investigations based on X-ray derived structures have focused on the roles of the substrate phosphate, ribofuranosyl and pyrimidine moieties in OMPDC’s catalytic proficiency ((*k*_cat_/*K*_m_)/*k*_non_).^23–26^ For instance, the R203A mutation in the phosphate-binding region increases *K*_m_ by approximately three orders of magnitude with minimal impact on *k*_cat_.^27^ Similarly, decarboxylation of the truncated substrate 1-(β-D-erythrofuranosyl)orotic acid (EO) in the presence of phosphite dianion shows a three-order-of-magnitude increase in *K*_m_, with little effect on *k*_cat_.^28^ These findings suggest that the phosphate group in OMP primarily enhances substrate binding by lowering *K*_m_, with minimal effect on the unimolecular reaction rate *k*_cat_.

Despite a significant reduction in activation enthalpy in the enzyme-catalyzed reaction compared to that in water, the Mt-OMPDC catalyzed decarboxylation still faces an enthalpic activation barrier of 15.5 kcal/mol after formation of the enzyme-substrate complex. Crystal structure comparison between the substrate-analog (SA) bound form and the TSA-bound form shows minimal structural differences, with a global RMSD of 0.3 Å (Fig. S1). The interaction distances between key residues and bound molecules differ by only about ±0.3 Å in the SA and TSA-bound forms, suggesting that the enzyme undergoes only slight structural change in the presence of a tight-binding inhibitor and arguing against "tight transition state binding" as the explanation of catalysis.

Theories of reaction rate in the condensed phase differ significantly from gas phase kinetics, ultimately being dependent on the structure and dynamics of the solvent. In particular, the kinetic and potential energy of the solvent must undergo dynamical transfer to the reactant(s), a process that takes place via transient environmental perturbations that create local changes in bond distances, angle and electrostatics required for barrier crossings.^29–31^ Extending these ideas to enzyme catalysis requires the incorporation of the protein scaffold together with its bound and surrounding water molecules into the dynamical solvation sphere.^32,33^ The first step in reaching transition state energies in enzymes is generally attributed to a conformational landscape; comprised of broadly distributed and rapidly inter- converting protein substates (ES) with energies for exchange close to the ambient temperature (∼0.6 kcal/mol at RT). The much larger experimental enthalpic barrier for Mt-OMPDC of 15.5 kcal/mol raises the question of how such a large energy gap can be overcome to create the millisecond time scale reactivity of both OMPDC and the majority of thermally activated enzymes.^6,34^

Temperature-dependent hydrogen-deuterium exchange mass spectrometry (TDHDX), when coupled with site-specific mutation that impairs the activation energy of catalysis (*E*_a_(*k*_cat_)), serves as a powerful tool for identifying spatially resolved protein networks, connecting the enzyme’s active site to the solvent bath. These networks are proposed to undergo thermal activation through collisions with solvent, leading to rapid structural and vibrational changes that modulate the active site environment and facilitate barrier crossings of the enzyme-substrate (ES) complex.^33,35,36^ This behavior offers a fundamental mechanism by which enzymes integrate physical processes within their scaffold and solvent interactions to drive chemical transformations at secluded active sites.^36^ TDHDX has been used across various enzyme systems, such as soybean lipoxygenase,^37,38^ dihydrofolate reductase,^39^ murine-adenosine deaminase,^40,41^ enolase,^42^ catechol O-methyl transferase,^43,44^ to identify reaction-specific networks for thermal energy transmission from solvent to active site. In this study, we report a detailed analysis of TDHDX on a ligand-bound form of wild-type (WT) Mt-OMPDC and a mutant that impairs the activation enthalpy, identifying four spatially resolved networks in the enzyme that connect their respective protein/water interfaces to key positions proximal to the bound substrate.

## Results

*Kinetic Analyses of Wild-Type and Mutant Forms of Mt-OMPDC*: WT and mutant forms of Mt- OMPDC were expressed and purified following established protocols^5^ with modifications (SI, Section- 2). The mutational strategy introduced subtle packing defects by replacing a hydrophobic residue with hydrophobic variants of altered shape and volume. This approach enables us to modify protein flexibility/motions without a significant alteration of electrostatics within the active site. The goal is to evaluate possible correlations between introduced changes in protein flexibility and the primary kinetic parameters *k*_cat_ and *K*_m_ together with the activation energy for *k*_cat_, *E*_a_(*k*_cat_). Enzyme kinetics were performed by spectroscopic monitoring of the product formation (SI, Section-3). A range of mutants were generated to identify the most effective function-impairing mutation, i.e., defined as one that minimally impacts *k*_cat_ (where *k*_cat_ is limited by rate of chemical step) and *K*_m_ but significantly alters *E*_a_(*k*_cat_). In the case of Mt-OMPDC, we have targeted residues forming a hydrophobic patch (Fig. 1c) within Van der Waals contact of substrate and with each other.

Notably, hydrophobic mutations at Leu123 (L123A, L123I, L123V) lead to very modest changes in *k*_cat_ and *K*_m_ (2-3 fold) with a 2.2 kcal/mol increase in the *E*_a_(*k*_cat_), Table 1 and Fig. S2. The rise in activation energy shows a linear correlation with the hydrophobicity of the substituted residue with the exception of the Gly substitution that resembles WT (Fig. 1e). The correlation between *k*_cat_ and hydrophobicity is less pronounced (Fig. S3). In the course of these experiments we interrogated other single site hydrophobic side chain alterations, such as Ile68, Ile96, and Val155 (Fig. S4), which in all instances led to small changes in *E*_a_(*k*_cat_) (ca. ±1 kcal/mol) along with decreases in *k*_cat_ that for the majority (>80%) were no more than four-fold, Table S1. The generally minor impact on kinetic parameters at these sites shows that the structure of Mt-OMPDC is robust and relatively insensitive to random mutations within hydrophobic side chains throughout the core. Residue L123 is part of the hydrophobic patch near substrate, situated at the start of loop5 and within Van der Waals distance of the K72 residue. Based on the presented kinetic analyses, WT and the L123A variant of Mt-OMPDC were selected for an in-depth investigation using single-temperature HDX followed by TDHDX.

**Table 1:** Kinetic parameters of wild-type and L123 mutants of Mt-OMPDC (with standard deviation from triplicate measurements in parentheses)

| | $k_{\text{cat}}$ ($\text{s}^{-1}$) at 25 °C | $K_m$ ($\mu\text{M}$) at 25 °C | $E_a(k_{\text{cat}})$ (kcal/mol) |
| --- | --- | --- | --- |
| WT | 4.3(0.4) | 1.4(0.2) | 15.5 (0.3) |
| L123A | 1.4(0.1) | 3.1(0.3) | 17.7(0.3) |
| L123V | 2.3(0.2) | 1.4(0.3) | 16.1(0.5) |
| L123I | 2.6(0.1) | 1.1(0.1) | 16.7(0.5) |
| L123G | 3.6(0.1) | 2.2(0.2) | 15.1(0.3) |

*Single-Temperature HDX Analysis*: Single-temperature HDX has been widely used to gain insights into protein-ligand, and protein-protein interactions.^45,46^ HDX measurements at 35°C were analyzed for Mt-OMPDC, to assess changes in deuterium uptake due to mutation and ligand binding. Detailed protocols for the HDX and data collection are provided in the SI, Section-4. Briefly, WT(apo) and L123A(apo) were directly incubated with deuterated buffer, while WT(L) and L123A(L) were pre-incubated with saturating concentrations of the tight-binding inhibitor 6-azaUMP (Table S2), introduced as a TSA^10,11^, for 30 minutes prior to exposure to D_2_O. The HDX experiments were conducted across 14 timepoints (10 seconds to 4 hours) followed by quenching, proteolytic digestion and liquid chromatography-mass spectrometry (LC-MS) analysis. The D-uptake was analyzed using 19 non-overlapping peptides providing 94% protein sequence coverage (Fig. S5 and Table S3). Experiments were conducted using biological replicates and the EX-2 condition for HDX was verified by inspection of time-dependent mass envelopes, and data were corrected for back-exchange (Table S4).

Fig. S6 shows comparative, time-dependent D-uptake for peptides from both apo- and ligand-bound forms of WT and L123A at 35°C. The variant L123A(apo) is seen to be greatly destabilized in the region at or close to the mutational site (peptides 122-133 and 133-141). Additionally, two peptides at the dimer interface (94-110 and 71-88) show a marked increase in HDX after 20 minutes, suggesting increased dissociation of the dimeric form of L123A(apo). The implied destabilization of L123A(apo) is consistent with *T*_m_ measurements that indicate a *T*_m_ of 60°C compared to *T*_m_ of 75°C for WT(apo) (Fig. S7). With this degree of instability, L123A(apo) was concluded to be a poor candidate for comparative HDX studies across a wide temperature range.

By contrast, the comparative HDX analysis of L123A and WT in the presence of 6-azaUMP is seen to eliminate the extreme effects observed for apo-protein, bringing the behavior of L123A(L) at 35°C closer to WT(L) for virtually every peptide analyzed (Fig. S6). These observations are supported by an observed 15°C increase in the *T*_m_ of L123A(L) to 74°C, that can be compared to a *T*_m_ of 81°C for WT(L) (Fig. S7). On this basis, it was decided to restrict detailed temperature variable HDX comparisons of L123A to WT to the protein-ligand complexes (see *Multiple Temperature TDHDX Analysis of WT(L) and L123A(L) below*).

*Quantitative Analyses of the Impact of Mutation and Ligand Binding on HDX at 35°C.* Differences between D-uptake for L123A and WT as well as the impacts of ligand binding were assessed by estimating the normalized percentage change in D-uptake (Δ*D*(%)) using equation 1:

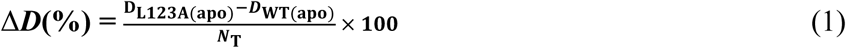

where *D*_L123A(apo)_ and *D*_WT(apo)_ represent the D-uptake within a given peptide in L123A(apo) and WT(apo), respectively, at given time. *N*_T_ is the total number of exchangeable amides in that peptide. The impact of ligand binding on enzyme (both WT and L123A) was estimated by comparing *D*_E(L)_ and *D*_E(apo)_, which represent the D-uptake within a specific peptide in the presence or absence of ligand, respectively. Representative Δ*D*(%) values between L123A(apo) and WT(apo) after 10 minutes (intermediate time regime for HDX) and 120 minutes (slow time regime for HDX) are summarized in Table S5 and mapped onto the crystal structure for peptides with Δ*D*(%) at a 3*σ* or higher level of significance in Fig. 2a-b; visualization is represented by blue-white-red heatmaps, showing lower (blue) or higher (red) D-uptake, in mutant, respectively. An increase in protein flexibility for L123A(apo) near the site of mutation is seen at 10 minutes (Fig. 2a) in peptides 122-133 and 133-141, along with some decreased flexibility around the periphery of the protein. While the regions of protein coverage that conform to 3*σ* differences are somewhat different at the 120 minutes incubation (Fig. 2b), a clear-cut trend of increased exposure at longer times is observed for L123A-derived peptides near the dimer interface (peptides 94-110, 111-121 and 71-88). These results corroborate the conclusions from *T*_m_ measurements of reduced stability for L123A, with the pattern of changes in HDX consistent with reduced stability at the dimer interface.

**Figure 2.**
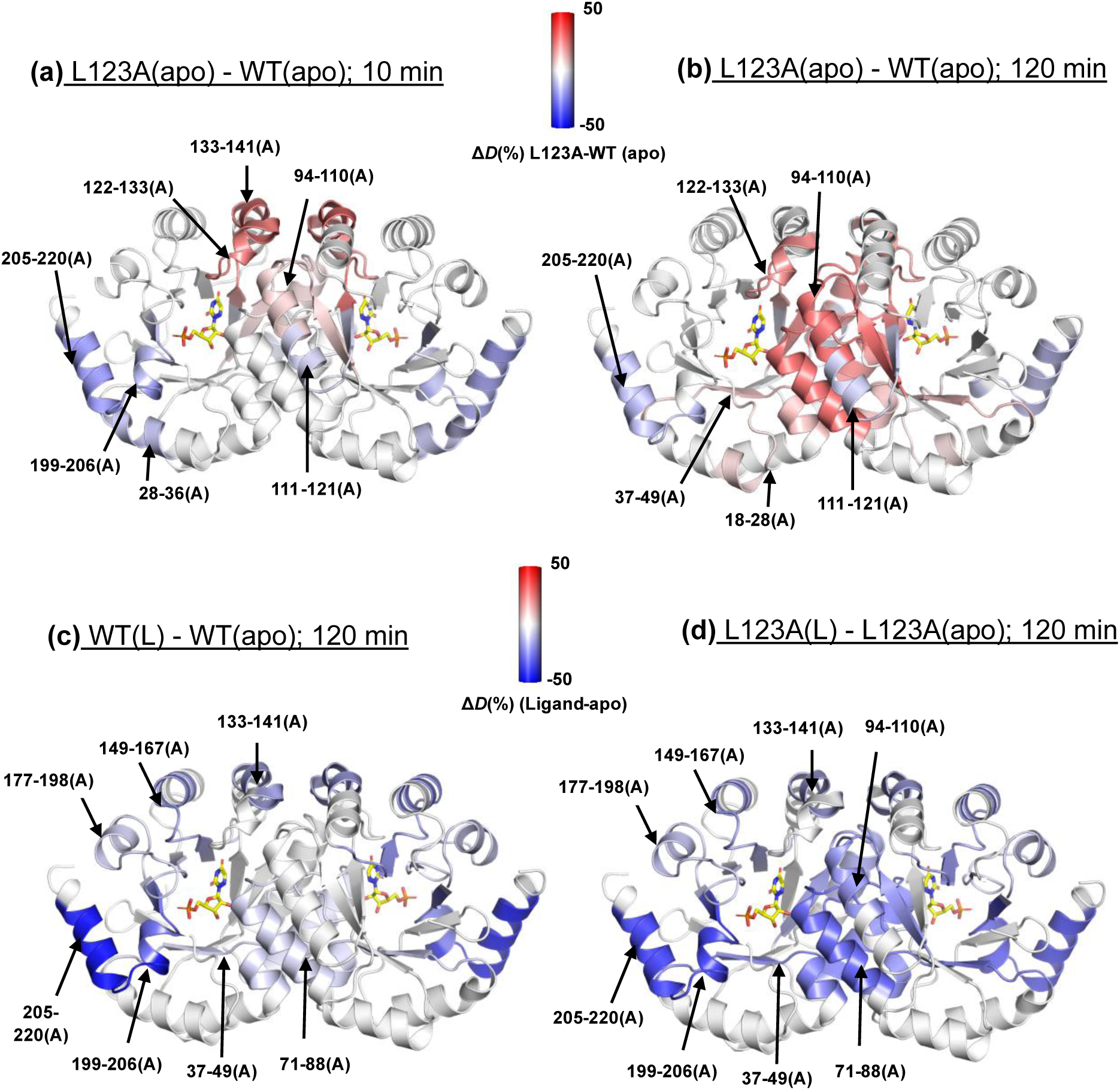
Single-Temperature HDX Analysis at 35 °C to Assess the Impact of Mutation and Ligand Binding on Thermal Stability of Mt-OMPDC. Panels (a) and (b) show the effects of the L123A mutation in the apo form, with panel (a) representing normalized percentage change in D-uptake, Δ*D*(%), after 10 minutes of HDX and panel (b) after 120 minutes. The Δ*D*(%), calculated using Equation 1 (see text), quantifies differences in D-uptake for each peptide due to the mutation. These changes were mapped onto the Mt-OMPDC crystal structure (PDB: 3G1A) using a blue-white-red gradient, where blue and red indicate decreased and increased D-uptake, respectively, in the mutant (L123A) compared to the wild type (WT). Only changes greater than 0.5 Da and 3*σ*, were considered, and peptides showing significant changes are marked with arrows only on the A monomer to avoid overcrowding. Panels (c) and (d) display the effects of ligand binding on WT and L123A after 120 min of HDX. These Δ*D*(%) changes were similarly mapped using the blue-white-red gradient, with blue indicating decreased D-uptake (enhanced structuring) and red indicating increased D-uptake (reduced structuring) in the ligand-bound form compared to the apo form.

The direct impact of ligand binding on protection against HDX at a 3*s* level of significance is also very informative in this regard (see Table S6 for the complete tabulation of data for WT and L123A). In Fig 2c-d, we focus on HDX changes at 120 minutes for WT(L)-WT(Apo) and L123A(L)-L123A(apo). In both enzyme forms, there is a similar degree of protection from binding of ligand at the periphery of the protein. By contrast, there is a marked increase in protection seen for L123A(L) - L123(apo) at the dimer interface peptides 94-110 and 71-88. Despite the absence of definitive 3*s* data for comparison to peptide 94-110 in WT(L)-WT(apo), protection by 6-azaUMP against structural disruption at the dimer interface in L123A(apo) is very apparent from the behavior of peptide 71-88.

*Comparative HDX Patterns for L123A(L) vs. WT(L) at single temperatures.* The TSA-bound form of WT reflects the conformational dynamics of the catalytically relevant closed states of OMPDC. When HDX differences between L123A(L) and WT(L) were analyzed at 35°C (Tables S7A-B, Fig. S8a-b), most peptides showed minimal changes. In the case of peptide 37-49, there is a slight increase at 10 minutes that disappears after two hours, and peptides 60-71 and 122-133 indicate ∼10% increase at two hours. Similar trends were observed at 50°C (near *T*_opt_ = 65°C), with only minor differences after 10 minutes and two hours (Tables S7C-D, Fig. S8c-d). This analysis highlights the limitation of comparative single-temperature HDX in capturing changes to conformational dynamics from site-specific mutation. The use of TDHDX provides a far more insightful connection between protein scaffold motion and enzyme activity.

*TDHDX Analysis of WT(L) and L123A(L)*: The goal of TDHDX is to determine local, site-specific changes in activation energies, *E*_a_(*k*_HDX_), for the protein scaffold of a chosen mutant that can be directly compared to the impact of mutation on *E*_a_(*k*_cat_). Sample preparation and HDX timepoints used in TDHDX analysis are the same as the single-temperature HDX, extended to seven temperatures (SI, Section-4). The D-uptake as function of time for peptide is analyzed using a three-exponential equation,

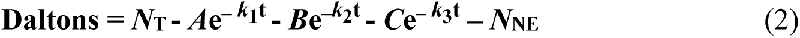

where *k*_1_*, k*_2_, and *k*_3_ define the fast, intermediate and slow HD exchange rate regimes, respectively, with *A*, *B*, and *C* indicating their respective amplitudes. *N*_T_ represents the total number of exchanging amides, and *N*_NE_ the non-exchanging amides. Boundary conditions for the different HDX exchange regimes were determined by initially fitting the D-uptake vs. ln(*T*) data using standard time regimes.^42^ Subsequent manual curation of these exponential fits was used to refine the time-dependent boundaries for slow, intermediate and fast exchange. The peptide-specific fitting parameters were determined as detailed in SI Section-5, and summarized in Tables S8–S9. The final fitting parameters obtained for WT(L) and L123A(L) are provided in Tables S10–S13. Due to time constraints in manual quenching of HDX, precise measurement of *k*_1_ is not feasible, and these studies are focused on the measurable exchange time regimes, *k*_2_ and *k*_3_. The rate constant for HDX, *k*_HDX_, is generally well determined using a weighted average rate constant^40,41,42^ (equation 3), although in some instances the data may be better analyzed using the temperature dependence of the individual rate constants.^40^

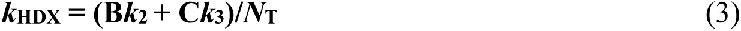

Temperature-dependence of *k*_HDX_ was assessed by performing HDX at seven temperatures (between 15-55°C) for both WT(L) and L123A(L). Arrhenius plot analysis of *k*_HDX_ yields *E*_a_(*k*_HDX_) = Δ*H*^⧧^(*k*_HDX_) + RT. Under EX-2 conditions (*k*_closed_ *>> k*_int_), Δ*H*^⧧^(*k*_HDX_) is comprised of the sum of Δ*H*°_open_ (enthalpy of opening-closing equilibrium; *K*_open_) and Δ*H*^⧧^_int_ (activation enthalpy of the intrinsic HDX rate constant, *k*_int_). Since *k*_int_ is typically unaffected by a single-site mutation (with the possible exception of the mutation-containing peptide), subtraction of Δ*H*^⧧^(*k*_HDX_) for WT from L123A leads to a direct determination of the impact of mutation on Δ*H*° for *K*_open_.

In this manner, time-averaged TDHDX analysis of WT and the *E*_a_(*k*_cat_)-impaired mutant (L123A) provides a probe for the sought-after relationship between mutation-induced changes in local scaffold flexibility (Δ*H*°) and activation energies for the chemistry at the active site (*E*_a_*(k*_cat_)). The full set of time and temperature dependencies of D-uptake in WT(L) and L123A(L) can be found in Fig. S9. The full set of comparative Arrhenius plots are presented in Fig. S10 and *E*_a_(*k*_HDX_) values for each peptide are listed in Table S14. In almost every instance, fittings that used weighted average rate constants or individual *k_2_* values yielded comparable results. Out of the 19 peptides, 16 showed significant values for *E*_a_(*k*_HDX_), while the remaining 3 did not exhibit an Arrhenius dependence of the rate. Among these 16 peptides, 8 yielded statistically significant (2*σ*) values for Δ*E*_a_(*k*_HDX_).

*Illustrative Comparison of TDHDX of L123A(L) to WT(L) for Peptides 37-49 and 94-110*. Figs. 3a- b display HDX data in WT(L) and L123A(L) as a function of temperature for peptide 37-49 (representing substrate ribose-binding region). Fig. 3c shows the Arrhenius plot for this peptide, with *E*_a_(*k*_HDX_) increasing from 9.0 ± 0.9 kcal/mol in WT(L) to 15.9 ± 1.2 kcal/mol in L123A(L). The Δ*E*_a_(*k*_HDX_) of 6.9 ± 1.1 kcal/mol, calculated by subtracting the *E*_a_(*k*_HDX_) of WT(L) from L123A(L), indicates that mutation increases protein’s rigidity in this region. Similarly, Figs. 3d-e show the D- uptake traces for peptide 94-110 (representing the dimer interface), with the Arrhenius plots presented in Fig. 3f. The *E*_a_(*k*_HDX_) is 12.3 ± 1.3 kcal/mol for WT(L) and 13.5 ± 1.2 kcal/mol for L123A(L), leading to a statistically insignificant Δ*E*_a_(*k*_HDX_), confirming no impact of mutation on local unfolding dynamics at the dimer interface when ligand is present.

**Figure 3.**
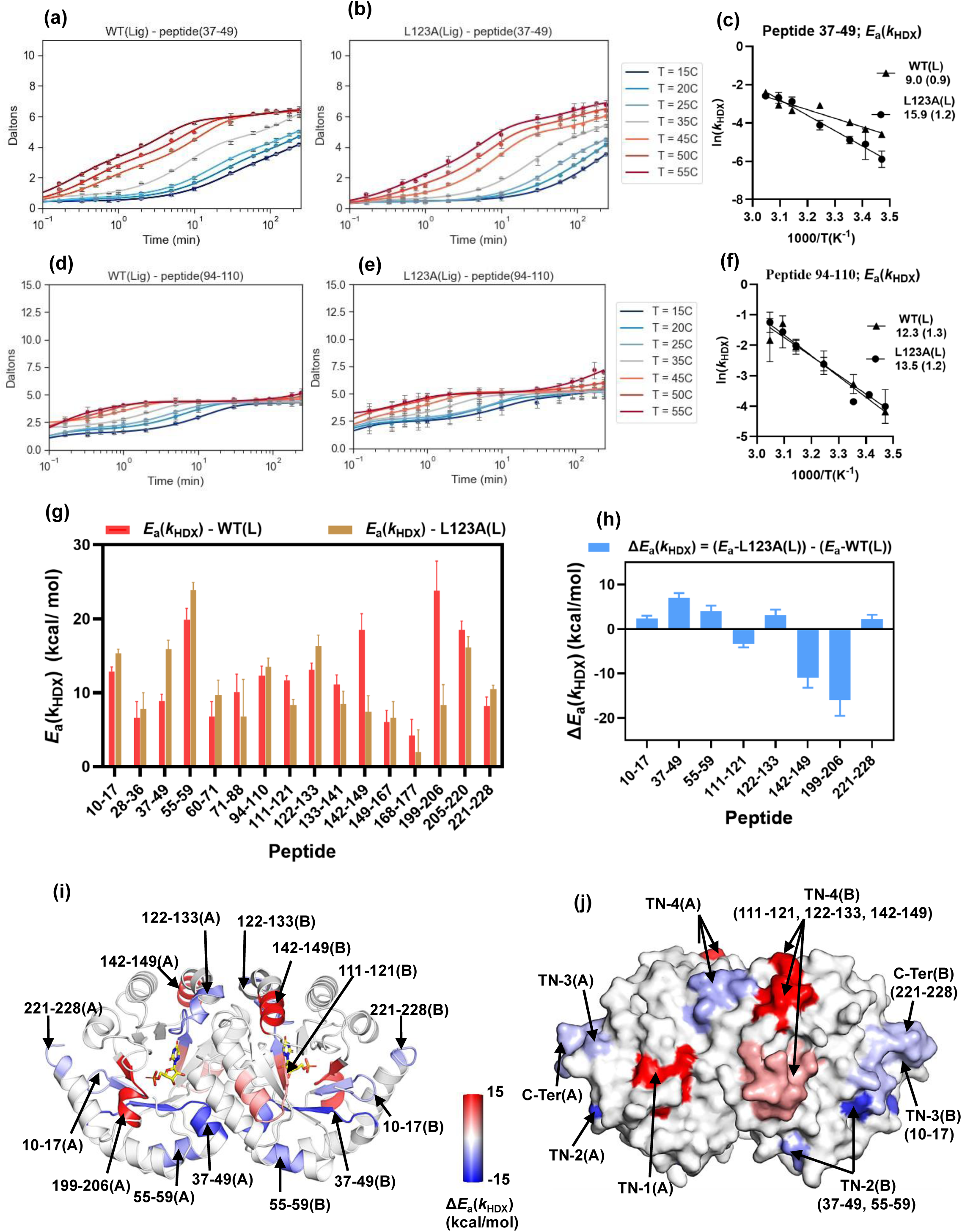
Temperature-Dependent HDX-MS Analysis of WT(L) and the *E*a(*k*cat) Impairing-Mutant L123A(L) Variants of Mt-OMPDC. Panels (a) and (b) D-uptake as a function of time and temperature for peptide 37-44 (representing ribose-binding motif) in the TSA-bound forms of native WT(L) and mutant L123A(L), respectively. (c) Arrhenius plot of ln(*k*HDX) vs 1000/T for peptide 37-44. Panels (d) and (e) D-uptake as a function of time and temperature for peptide 94-110 (representing dimer interface) in the TSA-bound forms of WT(L) and L123A(L), respectively. (f) Arrhenius plot of ln(*k*HDX) vs 1000/T for peptide 94-110. (g) Bar graph representing the HDX activation energy, *E*a(*k*HDX), values of peptides in WT(L) and L123A(L). (h) Bar graph representing Δ*E*a(*k*HDX), calculated as *E*a(*k*HDX) for (L123A(L)) minus *E*a(*k*HDX) for WT(L)). To ensure statistical significance, only Δ*E*a(*k*HDX) values greater than 2*σ*. (i) Cartoon representation of the Mt-OMPDC homodimer, highlighting peptides with significant Δ*E*a(*k*HDX). Regions with changes in Δ*E*a(*k*HDX) are shown using a red-white-blue gradient, with blue indicating positive Δ*E*a(*k*HDX) and increased rigidity in mutant, and red indicating negative Δ*E*a(*k*HDX) and increased flexibility in L123A(L) relative to WT(L). Peptides with significant changes in Δ*E*a(*k*HDX) are marked with arrows. (j) The space-filled model highlights regions with Δ*E*a(*k*HDX) changes using a red-white-blue gradient, revealing dynamic thermal networks (TN) connecting solvent-exposed regions to the active site. Thermal Network 1 (TN-1) corresponds to the phosphate-binding loop. Thermal Network 2 (TN-2) represents the sugar-binding region. Thermal Network 3 (TN-3) links TN-1 and TN-2 to facilitate synergistic interactions. Thermal Network 4 (TN-4) represents the catalytic loop. It is worth noting that TN-1 is present in both monomers but only shown in one monomer (TN-1 in A) as is on backside of surface the monomers B.

*Distributed Differences in ΔE*_a_(*k*_HDX_) *for WT(L) and L123A(L)*. The 16 peptides with measurable *E*_a_(*k*_HDX_) values are summarized in Fig. 3g and the Δ*E*_a_(*k*_HDX_) values for 8 peptides exceeding 2*σ* are shown in Fig. 3h. These peptides are mapped onto the structure of Mt-OMPDC using a blue-white-red gradient heatmap (Fig. 3i), highlighting spatially resolved regions with altered enthalpy of local unfolding due to the *E*_a_(*k*_cat_)-impairing mutant. In this heatmap, blue indicates increased rigidity in the mutant relative to WT, while red indicates increased flexibility. Table 2 summarizes the secondary structure composition and key catalytic residues for peptides with significant changes.

**Table 2.** HDX activation energy (*E*a(*k*HDX)) for WT and L123A mutant Mt-OMPDC, highlighting selected peptides along with their structural features and key catalytic residues.

| Peptide | $E_a(k_{\text{HDX}})$; WT(L) | $E_a(k_{\text{HDX}})$; L123A(L) | $\Delta E_a(k_{\text{HDX}})$; L123A(L)-WT(L) | Properties |
| --- | --- | --- | --- | --- |
| 10-17 (TN-3) | 12.9 (0.6) | 15.3 (0.6) | 2.4 (0.6) | The N-terminal $\beta 1$ of TIM-barrel forms H-bonds with $\beta 8$ (199-206; TN-1) and $\beta 2$ (37-49; TN-2) and interacts via van der Waals forces with the C-terminal peptide (221-228). |
| 37-49 (TN-2) | 9.0 (0.9) | 15.9 (1.2) | 6.9 (1.1) | This peptide includes the K42 ribose-binding residue, with $\beta 2$ strand forming H-bonds with $\beta 1$ (10-17; TN-3) and engaging in van der Waals interactions with peptide 55-59. |
| 55-59 (TN-2) | 19.9 (1.4) | 23.8 (1.0) | 4.0 (1.3) | A solvent-exposed peptide containing $\alpha 2$ of the TIM-barrel interacts via van der Waals forces with peptide 37-49. |
| 111-121 (TN-4) | 11.7 (0.6) | 8.3 (0.8) | -3.4 (0.7) | Contains the $\alpha 4$-loop-$\beta 5$ region of the TIM-barrel and neighbours the mutation site in peptide 122-133. |
| 122-133 (TN-4) | 13.1 (0.9) | 16.3 (1.5) | 3.2 (1.3) | Contains the loop5 of Mt-OMPDC, including the mutation site L123 and the intermediate-stabilizing residue S127, and forms van der Waals contacts with peptides 111-121 and 142-149. |
| 142-149 (TN-4) | 18.5 (2.4) | 7.4 (2.2) | -11.1(2.2) | Contains the $\alpha 5$ region of the TIM barrel and includes I142, which interacts with L122 (peptide 122-133) from the loop5 and F134' residue in other monomer. |
| 199-206 (TN-1) | 23.9 (4.0) | 8.4 (2.8) | -15.5 (3.5) | Phosphate-binding region ($\beta 8$-loop8) containing R203 and forms H-bonds with $\beta 1$ (10-17; TN-3). |
| 221-228 (C-terminus) | 8.1 (1.2) | 10.5 (0.5) | 2.3(0.9) | Contains the solvent-exposed C-terminus, forming van der Waals contacts with the TN-3 (10-17), which connects TN-1 and TN-2, thereby linking the C-terminus to the active site. |

Peptide 199-206, containing the R203 side-chain that interacts with the phosphate group of the substrate (phosphate-binding region), shows the largest reduction in *E*_a_(*k*_HDX_) (-15.5 kcal/mol) implying significantly increased flexibility in this region due to mutation. While mutations within the phosphate-binding region typically raise *K*_m_ for substrate by 2-3 orders of magnitude,^27^ the L123A mutation causes only a 2-fold increase, suggesting minimal impact of this mutation on the ground state ensemble of ES complexes. By contrast, the observed impact of L123A on *E*_a_(*k*_HDX_) within this region indicates a mutation-induced disruption of protein packing that is expected to reduce the level of efficient thermal energy transfer from solvent to bound substrate that is intrinsic to the WT enzyme. Peptide 199-206 is seen to be exposed to the solvent surface, connecting the solvent bath to the active site (Fig. 3j) and is, thus, identified as thermal network 1 (TN-1) in Mt-OMPDC.

The structurally adjacent peptide 10-17 at the N-terminus, shows a small decrease in flexibility, Δ*E*_a_(*k*_HDX_) = 2.6 kcal/mol, as does the C-terminal peptide 221-228, Δ*E*_a_(*k*_HDX_) = 2.3 kcal/mol. The β1- strand in peptide 10-17 forms H-bonds with the β8-strand in peptide 199-206 (TN-1), creating a structural link between these regions. Additionally, peptide 10-17 engages in H-bonding with the C- terminal peptide 221-228, thus establishing a connection between TN-1 and the solvent-exposed regions at the N and C- termini of the protein (Fig. 4a).

**Figure 4.**
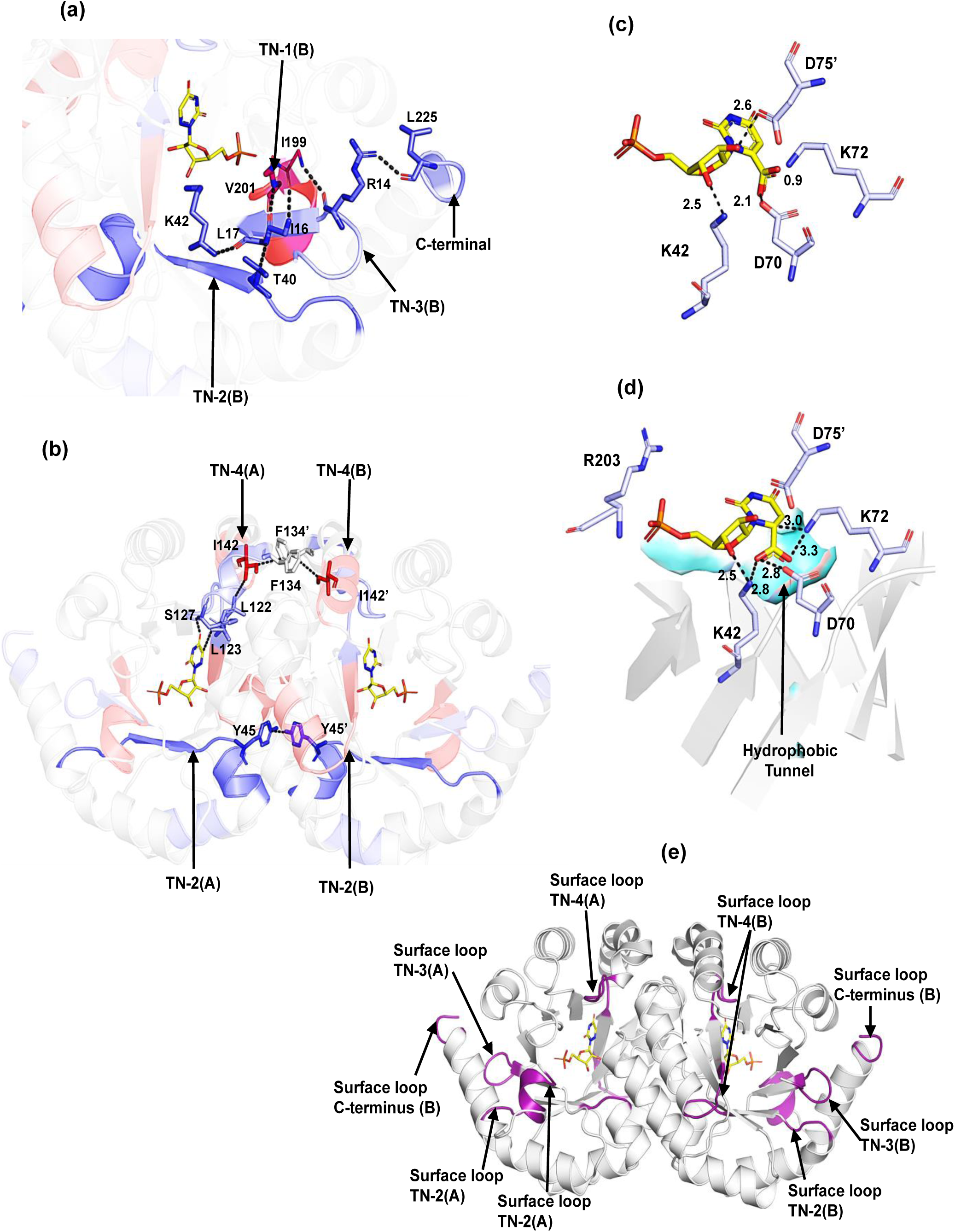
Synergistic Action of Intra- and Inter-Subunit Thermal Networks Drives Thermal Activation of Catalysis and Barrier Crossing in Mt-OMPDC. (a) Intra-subunit interactions between thermal-networks: Peptide 10-17 (TN-3) bridges TN-1 and TN-2 through H-bonding, enabling synergistic action. Hydrogen bonding between R14 and L225 connects the N- and C-termini. (b) Inter-subunit interactions: F134-I142’ (linking TN-4 and TN-4’) and Y45-Y45’ (connecting TN-2 and TN-2’) facilitates coordinated action of thermal networks between both subunits, alongside intra-subunit connectivity of TN-2 and TN-4. (c) The active site of OMPDC was modeled using the 6-azaUMP bound crystal structure (PDB: 3G1A). To represent the substrate- bound active site, OMP was positioned in place of 6-azaUMP in the structure. In this initial configuration, where the carboxylate group is in the plane of the pyrimidine group, it sterically clashes with the catalytic residue K72 (0.9 Å) and experiences electrostatic repulsion from D70 (2.1 Å). (d) Modeled active site of OMPDC, where the carboxylate group was bent out of the pyrimidine plane to avoid steric repulsion, positioning it into a hydrophobic patch. A minimum bend angle of 55° was required in this model to eliminate steric and electrostatic interactions. (e) Cartoon representation highlighting the spatial distribution of loop elements (purple) within the identified thermal networks in Mt-OMPDC. Surface-exposed loops of the C- terminus, TN-3 (N-terminus) and TN-2 are interconnected. Loops associated with TN-1 are solvent-exposed but connect to TN-2 through internal interactions rather than surface interactions.

Peptide 37-49 contains K42, a key residue for substrate binding and catalysis, that forms H-bonds with the substrate ribose 3’OH, and D70. K59A mutagenesis in Sc-OMPDC, equivalent to K42 in Mt- OMPDC, decreases *k*_cat_ by 100-fold and increases *K*_m_ by 1000-fold.^47,48^ A Δ*E*_a_(*k*_HDX_) of 6.9 kcal/mol for peptide 37-49 following introduction of L123A indicates increased rigidity near the sugar-binding region. Decreased flexibility is expected to perturb the ability of enzyme to achieve activated-complex configurations in a manner analogous to the perturbation that arises from increased flexibility within the phosphate-binding region. Enhanced rigidity of K42 (37-49) is also likely to disrupt the dynamic positioning of D70, a key residue for catalysis via substrate destabilization, and this may be a major source of the increased value of *E*_a_(*k*_cat_) for L123A. Peptide 55-59 is in Van der Waals contact with peptide 37-49 and also shows increased rigidity (Δ*E*_a_(*k*_HDX_) = 4.0 kcal/mol). Though not directly interacting with the substrate, its dynamics may impact peptide 37-49 and indirectly influence the thermal initiation of bond cleavage. Space filling representation reveals that peptide 37-49 and 55-59 connect the solvent bath to active site (Fig. 3j) and are ascribed to thermal network 2 (TN-2) in Mt- OMPDC. Note that the marked rigidification of peptide 37-49 in L123A was apparent from single-temperature analysis (Fig. S6). The β1-strand in peptide 10-17 at the N-terminus, which interacts with peptide 199-206 (TN-1) and the C-terminus, also forms H-bonds with the β2-stand in peptide 37-49 (TN-2) (Fig. 4a). This establishes a connection between TN-2 and the exposed C- and N-termini, leading to the designation for peptide 10-17 as a separate thermal network 3 (TN-3) that coordinates the behavior of TN-1 and TN-2.

Peptide 122-133 includes loop5 and the mutational site L123, which is part of a hydrophobic cluster, a pathway for CO_2_ release, and includes S127 that is proposed to stabilize the vinyl carbanion/carbene intermediate.^18^ TDHDX shows a 3.2 kcal/mol increase in *E*_a_(*k*_HDX_) for L123A(L) relative to WT(L), indicating increased local rigidity that may impair thermally initiated vinyl intermediate stabilization and contribute to the elevated *E*_a_(*k*_cat_) for L123A. Peptide 142-149 is in Van der Waals contact with 122-133 in both monomers (Fig. 4b). Very significantly, this region shows a large decrease in Δ*E*_a_(*k*_HDX_) by -11.1 kcal/mol. A second region adjacent to L123, peptide 111-121, also shows a decrease in Δ*E*_a_(*k*_HDX_) by -3.4 kcal/mol. The increased flexibility of 142-149 and 111-121 is also expected to impair contributions of thermally activated dynamics within loop5. All three peptides are solvent-exposed (Fig 3J) and constitute thermal-network 4 (TN-4) in Mt-OMPDC.

## Discussion

*Impact of Temperature on Mt-OMPDC Stability*: The original goal of performing a full TDHDX comparative analysis of WT(apo) and L123A(apo) forms of Mt-OMPDC proved untenable, due to instability of the variant at elevated temperatures. This is supported by a large reduction in *T*_m_ for L123A(apo) to 60°C relative to a *T*_m_ of 75°C for WT(apo). As anticipated, addition of tight-binding 6- azaUMP elevated *T*_m_ for both WT(L) and L123A (L), to values of 74° and 81°C, respectively, where the highest temperature used for TDHDX was 55°C.

Comparisons of patterns of HDX at 35°C and 120 minutes for L123A(apo) and WT(apo) (Fig. 2b) indicate the regional loss of WT structure at the dimer interface of L123A, implicating an enhanced proclivity to dimer dissociation as a primary source of thermal instability in L123A. This is supported by the multiple temperature HDX analyses of apo forms (Fig. S11), where the L123A(apo) variant shows a continuous rise in D-uptake as a function of both time and temperature for the peptides representing the dimer interface (71-88, 94-110 and 142-149). By contrast, the WT(apo) shows none of this behavior, consistently plateauing in D-uptake with increased time and elevated temperature (Fig. S11). Significantly, thermally induced, functionally correlated changes in protein flexibility for L123A(L) relative to WT(L) are found to lie almost exclusively outside of the dimer interface, with the exception of peptide 142-149 (Fig 3i), implicating different origins of observed temperature responsive behavior for enzyme stability vs. function. A full understanding of the molecular origins of thermostability in WT Mt-OMPDC will require further investigation.

*Understanding the Temperature-Dependent Activation of OMPDC From the Perspective of Environmental Reorganization*: OMPDC achieves a ca. 17-order catalytic enhancement of the unimolecular decarboxylation of OMP, and is considered one of the most challenging enzyme reactions to fully rationalize. The literature indicates extensive analyses, with a focus on the contribution of interactions between specific residues and components of substrate on the enhancement of *k*_cat_ and the corresponding reduction in the free energy of activation, Δ*G*^‡^. ^49,50,51,52,53^ This focus on Δ*G*^‡^ derives from Pauling’s proposal that differential binding affinity of a transition state relative to substrate drives enzymatic reactions.^54^ However, X-ray crystallography reveals only slight conformational differences between the ES and ETS complexes of Mt-OMPDC (Fig. S1), indicating that static structures are unable to capture the physics that underlies the conversion of the ground state structure to its activated complex. ^55,56^

In this work, we turn our focus to the dramatic decrease in the Δ*H*^‡^ of the OMPDC reaction relative to its solution reaction (Fig.1a), further noting the dominant importance of reduced enthalpic barriers for the majority of enzyme-catalyzed reactions.^3,6,7^ Earlier studies have shown that studies of TDHDX, when performed with enzyme variants that display changes to *E*_a_(*k*_cat_), provide a spatial map of networks that participate in thermal energy transport from solvent to the reactive bond(s) of bound substrates.^37–43^ It is generally recognized that the catalytic competence of ES complexes will be critically dependent on the inherent flexibility of protein scaffolds that support a family of equilibrating protein substates with differing abilities to undergo barrier crossings. This property is most commonly attributed to a conformational landscape comprised of a high density of protein substates with small energetic differences (RT ∼ 0.6 kcal/mol). This formalism recognizes the inherent importance of the surrounding protein scaffold in accelerating reaction rates, while leaving open the question of the physical process whereby enzymes are able to overcome enthalpic barriers that are dominantly in the range of 10 kcal/mol.^3,6,7^ This value for enthalpic barriers is further elevated in the case of thermophilic enzyme, as seen in the case of Mt-OMPDC where Δ*H*^‡^ = 15.5 kcal/mol.

In condensed phase reactions, activated complex formation depends on diffusional encounters of reactants with each other together with subsequent alterations in solvent structure/dynamics that transfer energy to preorganized ground state encounter complexes. The role of environmental reorganization for chemical activation in the condensed phase has been formalized by Marcus theory, which recognizes *solvent reorganization (λ)* as the primary kinetic barrier, creating an environment complimentary to and capable of stabilizing changes in bond distances, bond angles and electrostatics that are prerequisites for functional barrier crossings.^29,30^ Marcus theory also differentiates environmental reorganization from the preorganization process that brings reactants into close proximity. In enzymatic reactions the latter is provided by a sequestered active site that is susceptible to sampling among a large number of equilibrating ES ground states.

Critically, the transitioning of this family of stable ground state ES complexes to activated complexes is distinctive, relying on high-energy structures that occur transiently and repeatedly within selective regions of the protein scaffold and its accompanying shell of bound water.^57,58^ Over the past decade, carefully designed experimental studies of a number of different enzyme systems have uncovered both

the spatial and temporal properties of such protein scaffold reorganization, formulating methods to filter out non-essential dynamics from functionally relevant protein motions that promote activated complex formation.^33,36^ The first step in such studies involves a spatial resolution of function-related protein motions, as afforded by TDHDX.

*Complex Thermal Networks Function in Mt-OMPDC*: From the TDHDX comparison of WT Mt- OMPDC to its *E*_a_(*k*_cat_) altering variant L123A, we have identified regions of protein that connect separate protein/solvent interfaces to key elements of the enzyme’s active site structure, Fig. 3i-j. Four distinct regions have emerged as spatially unique thermal networks (TNs): TN-1, terminating at the substrate phosphate-binding position, TN-2, terminating at the substrate ribose-binding region, TN-3 that connects TN-1 and TN-2, and TN-4 that includes loop5 and contains Ser127 that stabilizes the decarboxylated carbene/carbanion intermediate, Fig. 1d. Each is represented in Fig. 4a-b.

### TN-1

Substrate binding induces loop7 closure through a Q185/phosphate and additional interactions (Fig. S12), isolating the active site from solvent and achieving a catalytically competent state. Loop7 also interacts non-covalently with loop5 and the phosphate-binding region (R203). Although previous studies suggest that loop7 length affects the catalytic activation barrier in mesophilic vs. thermophilic forms,^5^ our TDHDX analysis of Mt-OMPDC does not identify loop7 as part of a thermal network. In an earlier TDHDX study of the enzyme enolase, the behavior of an active site loop closure was similarly differentiated from regions of the protein susceptible to mutation-induced changes in protein flexibility.^42^

Active site analysis indicates that the carboxylate group of OMP faces substantial electrostatic and steric interference from D70, with its proximity to K72 heightening this unfavorable interaction (Fig. 4c). To minimize these clashes, the carboxylate group is expected to bend out of the pyrimidine-ring plane, positioning itself in a nearby hydrophobic pocket that provides an ideal site for CO_2_ release,^52^ (cf. Fig. 4d). Similar out-of-plane bending is observed in crystal structures of OMPDC with other substrate analogs, suggesting a conserved mechanism that depends on effective substrate alignment to enhance catalytic activation.^19,20^ We propose that thermal activation in the phosphate-binding region (TN-1), modulated by R203-phosphate interactions modulates substrate positioning that transiently enhances the unfavorable energy arising from the interaction between the carboxylate and the D70- K72 pair. This, in turn, facilitates the out-of-plane bending of the carboxylate and forces it into the adjacent polar-hydrophobic interface (Fig. 4d), as one of the major driving forces for bond cleavage.

### TN-2

Similarly, thermal activation in the substrate ribose-binding region (TN-2) influences the K42- ribose ring interaction, thereby reinforcing the substrate’s carboxylate group dynamic positioning at a polar-hydrophobic interface. Additionally, K42 in TN-2 forms a hydrogen bond with D70 in the catalytic tetrad, with the expectation that TN-2 activation will adjust the positioning of D70, K72, and D75’ to fine-tune the active site’s electrostatic environment (Fig. 4d). The action of TN-1 and TN-2 in each monomeric unit of Mt-OMPDC is coordinated through their interaction with the β1-strand of peptide 10-17, assigned as TN-3, and also interacts with C-terminal peptide 221-228 that resides at a protein/water surface.

### TN-3

The presence of TN-3 (cf. Fig. 4a) is proposed to *synergistically link* activated protein motions within the phosphate (TN-1) and ribose (TN-2) regions of substrate, as a main driver of precise dynamical tuning of active site electrostatics. The resulting high energy, transient positioning of substrate facilitates the C-C bond cleavage reaction for release of CO_2_ into a hydrophobic pocket at a polar-hydrophobic surface, resulting in the formation of a stabilized carbanion-carbene intermediate (Fig. 1d).

### TN4

The carbene intermediate is stabilized by a H-bond from backbone amide of S127 in loop5 that is identified within TN-4. Thermal activation of TN-4 will modulate the H-bonding distance between the S127 N-H and O4 of the carbene, directly governing the stabilization energy of this catalytic intermediate (Fig. 4b). In Mt-OMPDC, hydrophobic residues from almost all β-strands (except β2) extend into the central barrel, forming a continuous hydrophobic tunnel (aqua-surface Fig. 4d). This tunnel, built from strategically arranged hydrophobic residues, establishes a network of isoleucine, leucine, and valine (ILV) side chains, which are involved in van der Waals contacts and identified through a contact-based structural unit algorithm^59^ (Fig. S13). The ILV network within the hydrophobic tunnel intricately connects all four identified thermal networks: TN-1 (containing I200), TN-3 (containing I16), and TN-4 (containing L123). Notably, TN-2 (containing K42) engages in van der Waals interactions with I16 of TN-3 and I200 of TN-1. Thus, the ILV network establishes a cohesive pathway that facilitates the synergistic action of all four thermal networks. In addition to intra-subunit connections between thermal networks, interactions such as F134-I142’ (connecting TN- 4 and TN-4’) and Y45-Y45’ (connecting TN-2 and TN-2’) may facilitate a synchronized action of thermal networks between subunits (Fig. 4b).

We conclude that the ability of OMPDC to reduce its enthalpic barrier for C-C bond cleavage by ca. 28 kcal/mol relative to the corresponding solution reaction (Fig. 1a) is a result of a protein scaffold that has evolved a preorganized active site in close proximity to multiple, interconnected thermal energy networks; the latter drive a thermally activated environmental reorganization process that creates transient complementary of the enzyme’s active site to the altered bond angles and electrostatics characteristic of substrate at the position of barrier crossing.

*Comparison of Thermal Networks in OMPDC to Other TIM Barrel Enzymes:* The protein networks identified in Mt-OMPDC provide a fourth example within the TIM barrel family of enzymes. The first distinctive feature of Mt-OMPDC is a mixed pattern of "more flexible/more rigid" in the regions of the loop5 and the substrate phosphate-binding region following insertion of the L123A (cf. Fig. 3i-j). A second feature is the large number of peptides, eight out of a total of 16 analyzed, that are altered in TDHDX by a single site substitution of Leu to Ala (Table 2). A third property is the detection of four distinctive thermal networks, differentiating OMPDC from the other TIM barrel enzymes thus far characterized by TDHDX. As summarized in Fig. 5 the TIM barrel enzymes murine adenosine deaminase, yeast enolase, and human catechol-O-methyltransferase, each reveal two thermal networks, separated by ca. 180°, 0° and 90°, respectively; in all cases, these converge at the reactive carbon center within the active site.^40–43^

**Figure 5.**
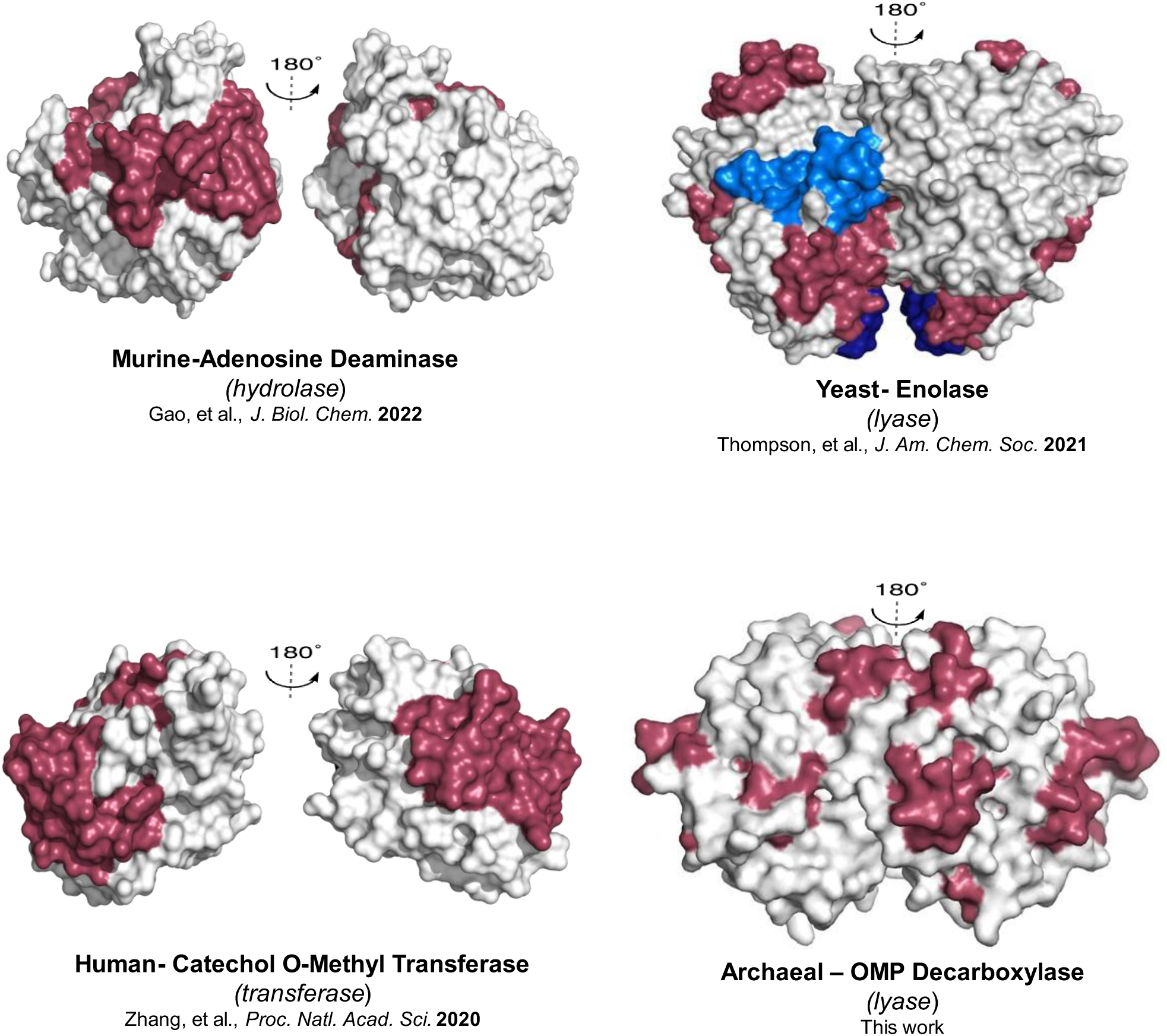
Thermal Energy Network Comparison Among TIM barrel enzymes. The final TDHDX-derived energy transfer pathways in murine adenosine deaminase (m-ADA), yeast enolase (Sc-ENO), and human catechol-O-methyltransferase (Hs-COMT) each consist of two distinct networks. In contrast, for archaeal OMP decarboxylase (Mt-OMPDC), which catalyzes one of biology’s most challenging reactions, four synergistic thermal energy networks have been identified. In every instance, these networks converge at the reactive bond of the substrate within the respective active site. The thermal energy network figure for m-ADA (Gao, et al., *J. Biol. Chem.* 2022), Sc-ENO (Thompson, et al., *J. Am. Chem. Soc.* 2021), and Hs-COMT (Zhang, et al., *Proc. Natl. Acad. Sci.* 2020) has been adapted from previously published studies, while the network for Mt-OMPDC is the result of this work.

We propose that the extensive thermally activated regions identified in Mt-OMPDC relative to the other TIM barrel enzymes studied is the result of enzyme adaptation to accommodate an enzyme reaction that requires such an enormous enthalpic reduction from the uncatalyzed reaction. The studied members of the TIM barrel family are further differentiated from the enzyme lipoxygenase, comprised of a different two domain structure.^32^ Lipoxygenases have been shown to catalyze a relatively "simple" C-H bond cleavage through quantum mechanical H-tunneling. In this instance, a single thermal energy network appears sufficient to achieve the synergistic reduction of the hydrogen donor-acceptor distance and alteration of Δ*G*^°^ that produce reactant and product energy wells supporting efficient wave function overlap.^38^

*Integrating Thermal Networks into Dynamic Models for Enzyme Activation:* A major mechanistic question is how motions at protein/solvent interfaces may propagate through enzyme networks to promote active site environments capable of rapid barrier crossings. Previous work from this lab on SLO^57^ and m-ADA^58^ extended TDHDX studies to include temporal probes of functionally relevant protein motions. In these instances, dynamic Stokes shift analyses, using fluorophores appended to the protein surface at spatially identified networks revealed a 1:1 relationship between the thermal activation barrier for reorganization at a protein/solvent interface (sub-ns to ps timescale) and the active site reorganization that transforms reactant to product on a slower ms scale. Time differences between the measured kinetic parameters are ascribed to different probabilities of a shared protein restructuring supporting the different reaction outcomes.

In SLO, its single thermal network facilitates long-range communication over ∼20 Å, channeling the entire activation energy for catalysis from solvent to active site. In m-ADA, enzymatic activation relies on more complex interactions from two networks originating on opposite faces of the protein structure and converging at the active site. In both instances, protein surface loops are identified as the dynamical feature that initiate rapid conformational shifts in response to interactions of bulk solvent with the protein/solvent interface. Loop dynamics have previously been shown to govern critical aspects of enzyme function, including substrate binding, active site desolvation, intermediate stabilization, turnover rates, specificity, allostery, and thermal adaptation between mesophilic enzymes and their psychrophilic and thermophilic homologs.^60–68^ In this context, Fig. 4e highlights the loops identified to be present at the termini of TN’s1-4 in Mt-OMPDC. Ongoing studies of Mt-OMPDC are focused on obtaining temporal resolution of ns-ps motions through Stokes shift studies of loop-specific chromophores, analogous to the approaches pursued with SLO^57^ and m-ADA.^58^

To conclude, while the importance of protein dynamics in enzyme function is well recognized, identifying functionally relevant motions amidst numerous other movements remains challenging. During the last decade, developed experimental methods have led to the identification of protein networks as a primary component in anisotropic energy transfer to effect productive reaction barrier crossings in enzyme-catalyzed reactions.^36,57,58^ Extending this approach to Mt-OMPDC, which catalyzes one of biology’s most challenging reactions, has uncovered an unusually complex network for thermal energy transfer. The results strongly suggest a correlation between the difficulty of the catalyzed reaction and the number of multiple site-specific conduits within the corresponding protein scaffold. *De novo* enzyme design, which has traditionally focused on refining active site interactions, has struggled to match the efficiency of natural enzymes^69,70^ due to the limited integration of dynamical components into design principles. Learning how to incorporate loop-based thermal energy transfer pathways within protein scaffolds offers a potentially powerful approach for the future creation of highly efficient biocatalysts.

## Acknowledgments

We thank Prof. Susan Miller, Prof. Nigel Richard, Dr. Gargi Jagdale, Dr. Kotchakorn Tsriwong, and Dr. Sabyasachi Sarkar for their helpful and stimulating discussions. We also thank Dr. J.P. Zaragoza and Dr. S. Gao for their valuable suggestions, and Dr. E. Thompson for providing the Python code used for data analysis. This research is supported by funding from the National Institutes of Health grants to J.P.K. (GM118117) and A.T.I. (1S10OD020062-01).

The supplementary file details experimental methods, structural comparisons, enzyme kinetics, single- and temperature-dependent HDX data on OMPDC, including figures, tables, and Python scripts for analysis.

## Author Contributions

P.D. and J.P.K. designed the research. P.D., A.S., J.L., and A.T.I. performed the research. P.D., A.S., and J.L. analyzed the data. J.P.K. conceived and supervised the project and acquired funding. P.D. and J.P.K. wrote the paper.

